# Effects of Classical Psychedelics on Implicit and Explicit Emotional Empathy and Cognitive Empathy: A Meta-analysis of MET task

**DOI:** 10.1101/2024.05.02.592231

**Authors:** Amit Olami, Leehe Peled-Avron

## Abstract

This meta-analysis investigates the effect of classic psychedelic drugs on empathy and focuses on cognitive and emotional empathy measured using the Multifaceted Empathy Test (MET). Empathy entails the ability to understand and share the feelings of another and is a significant component of social interaction. Several studies have examined the effects of psychedelic drugs such as LSD, psilocybin and ayahuasca on empathy, yet their overall effect has not been studied so far. In this meta analysis, we reviewed data from studies up to November 2023 with the aim of examining the effects of various psychedelic drugs on empathic abilities broadly. Our findings suggest that classical psychedelics significantly enhance explicit and implicit emotional empathy without affecting measures of cognitive empathy. The results emphasize the need to continue testing the therapeutic potential of classic psychedelic drugs.

## Introduction

Empathy is a complex psychological construct crucial for human interaction and communication [1,2]. It encompasses various dimensions, including cognitive and emotional empathy in implicit and explicit forms. Empathy is defined as the understanding of a person from their frame of reference rather than one’s own, or vicariously experiencing that person’s feelings, perceptions, and thoughts [3]. Empathy towards others can aid in creating more meaningful relationships and enhance our social interactions [4]. Cognitive empathy, also known as the Theory of Mind, involves the mental capacity to comprehend and process the thoughts and emotions of others, enabling us to grasp different perspectives and foster mutual understanding and respect in diverse interactions [5]. Emotional empathy, on the other hand, goes beyond mere understanding to an emotional resonance, allowing us to share in the joy, sorrow, excitement, or pain of others, and is crucial for building emotional bonds and providing comfort and support [6,7].

Emotional empathy can be further divided into implicit and explicit emotional empathy. Implicit empathy, often referred to as “arousal,” includes the automatic, unconscious aspect of emotional empathy in which the sharing of emotions arouses the observing individual. In contrast, explicit emotional empathy is a deliberate and conscious process requiring mental effort and conscious processing, enabling us to respond appropriately and empathetically toward other people’s emotions [3,8].

It is possible to change the levels of empathy with the help of various psychological and biological interventions [9-11]. One such intervention pertains to the use of classical psychedelics that can enhance empathic abilities. In recent years, researchers have discovered an interest in the effects of psychedelic drugs on social cognition in general and empathy in particular, and research has been done on the impact of these drugs on various types of empathy. Classical psychedelics, including Lysergic acid diethylamide (LSD), psilocybin, and ayahuasca, are all agonists at the 5-HT2A receptor, a serotonin receptor subtype [12,13]. These substances have shown potential in addressing various psychiatric conditions, including depression and anxiety [14-16], by breaking rigid mental patterns that are resistant to change [10,17]. LSD is derived from the ergot mushroom, leading to mood and perception changes at doses of 100-200 micrograms [18]. Psilocybin, derived from psilocybe mushrooms, affects the brain by converting it into psilocin, inducing similar shifts in perception and mood at doses of 20-25mg [19,20]. Ayahuasca, a South American brew containing N, N-Dimethyltryptamine (DMT) and Monoamine oxidase inhibitors (MAOIs), creates altered states of consciousness and vivid hallucinations [21,22]. Research presents diverse findings on psychedelics and empathy. LSD was demonstrated to enhance emotional empathy and sociality [23]. Similarly, psilocybin was found to increase emotional empathy [19,20] without affecting moral decision-making [20] and increasing well-being and creativity [19]. Additional research showed that ayahuasca increased cognitive empathy, emotional empathy, and well-being up to one week after the ingestion during a ceremony [10,11]. Based on our review of the professional literature, we have yet to find a meta-analysis that synthesizes this literature and examines the overall effect of classical psychedelic drugs on empathy. Therefore, our meta-analysis reviews recent studies in the field by synthesizing data from these studies on several types of empathy. These different forms of empathy collectively influence how we interact, communicate, and build relationships with others and how external factors, such as classical psychedelics use, can impact these forms of empathy, highlighting the complex interplay between psychological processes and biological mechanisms that underlie this social cognitive function [23].

In the present meta-analysis, we will examine the existing research literature on the effect of different classical psychedelics on a single task. We focused on the multifaceted empathy test (MET), measuring cognitive and emotional empathy using emotionally charged pictures. Meta-analyzing only one task across different studies leads to a more accurate assessment, allows for direct comparison of results, and provides deeper insights into specific factors associated with that task. It is especially important when comparing several classical psychedelics that are administered in different contexts with different doses and are measured at different times. We will examine the effect of LSD, psilocybin and ayahuasca on cognitive empathy and explicit and implicit emotional empathy to determine the effects of these classical psychedelics on various types of empathy.

## Methods

This study followed the PRISMA reporting guidelines for systematic reviews and meta-analysis [24].

### Search Strategy and inclusion criteria

From inception up to November 2023, a search for relevant articles was conducted in the PubMed (Medline), PsycINFO, and Scopus bibliographic databases using specific keywords: “LSD” OR “ayahuasca” OR “psychedelic” OR “lysergic acid diethylamide” OR “Psilocybin” AND “empathy” OR “emotion” OR “social cognition” OR “emotional empathy” OR “cognitive empathy” OR “implicit” OR “explicit” OR “implicit emotional empathy” OR “explicit emotional empathy”, AND “MET” OR “multifaceted empathy test”.

After a full-text review, if a study was deemed eligible, its reference list was manually scrutinized for other relevant studies.

Initially, duplicate records were eliminated, and subsequently, the titles and abstracts of the remaining publications were assessed. Any publications that did not meet the inclusion criteria were excluded. The remaining publications were further evaluated by thoroughly reviewing the full text. The inclusion criteria were then applied again. To be considered for inclusion, studies had to meet the following eligibility criteria: (1) they had to be original, peer-reviewed, full-text articles written in English; (2) they had to involve healthy human participants; (3) they had to include a single-dose or more of LSD or ayahuasca or psilocybin intervention; and (4) they had to include an outcome(s) related to the MET tasks, provided that they met the eligibility criteria mentioned above. Studies were excluded if they were reviews and/or meta-analyses, included non-human subjects, assessed previous classic psychedelics users retrospectively through self-report questionnaires, or were letters, comments, abstracts, or conference papers. There were no exclusions based on the age or sex of the subjects, classic psychedelics dose or frequency, administration routes, or study size (Figure 1).

**Figure 1.**
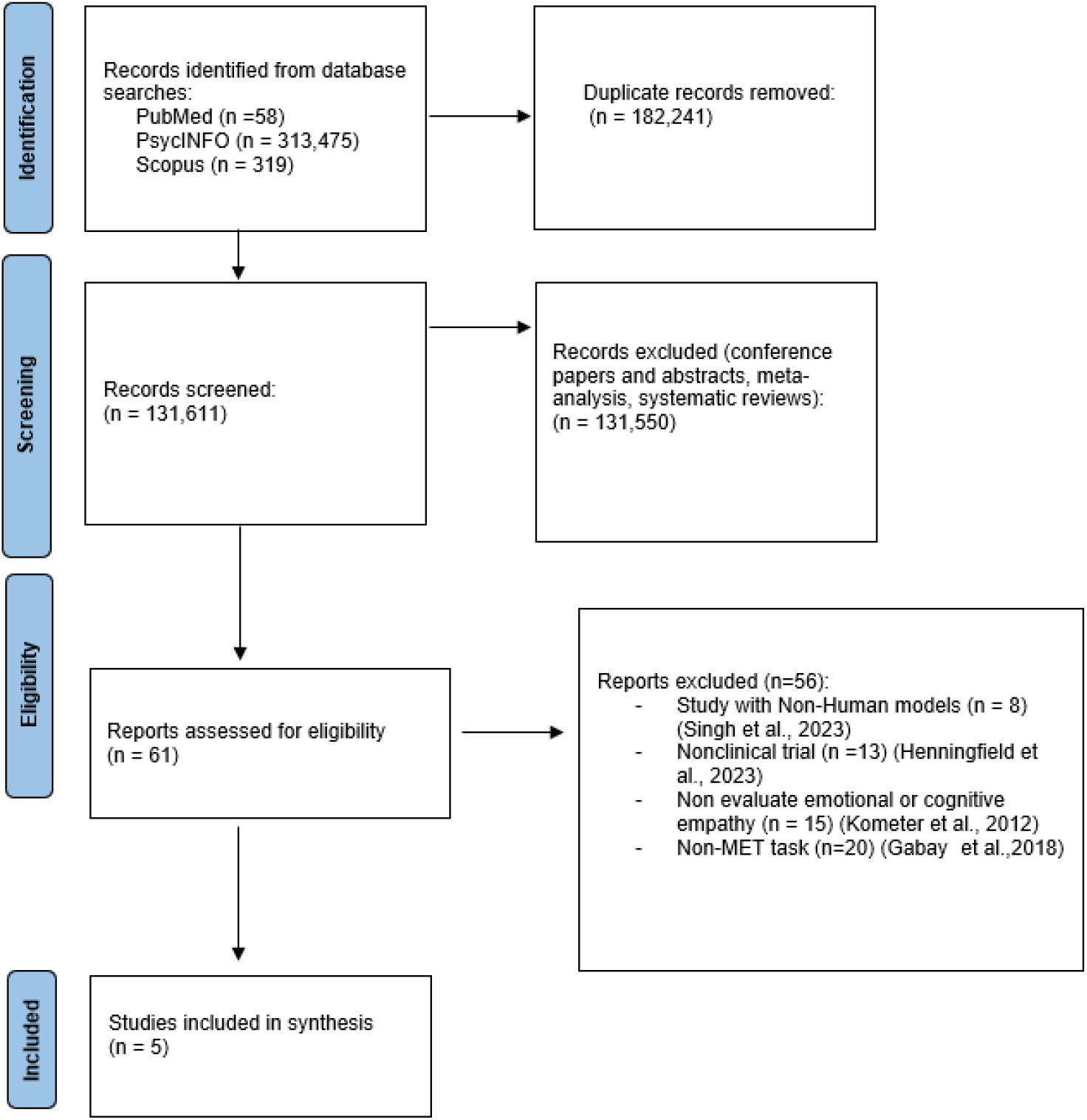
The search and selection procedure was used to identify studies for inclusion in this meta-analysis. Template provided by PRISMA (www.prisma-statement.org).

### Recorded measures and data extraction

The recorded variables included general information, such as the study’s authors, year of publication, number of subjects, and demographic variables, such as age, gender, and classic psychedelic dosage and kinds of substance (Table 1). For the calculation of effect sizes, means and standard deviations (SD) of the MET task results were collected. If this information was not available in the publication or provided by the authors, effect sizes were estimated using one of the following methods: (1) by extracting mean and SD from published figures (using PlotDigitizer software, available at http://plotdigitizer.sourceforge.net), (2) based on F values, or (3) based on p values.

**Table 1.** demographic details on the included studies.

| Authors and year | Sample size | M/F | Mean age + S.D | Substance | Dosage |
| --- | --- | --- | --- | --- | --- |
| Dolder et al, 2016 | 22 | 12M, 12F | 33 ± 11 | LSD | 100µg |
| Dolder et al, 2016 | 22 | 12M, 12F | 33 ± 11 | LSD | 200µg |
| Thomas Pokorny et al, 2017 | 32 | 17M,15F | 26.72 ± 5.34 | Psilocybin | 0.215mg |
| Mason et al., 2019 | 55 | 29M, 26F | 34.8 ± 8.9 | Psilocybin | 1.9mg |
| Uthaug, M et al, 2021 | 30 | 12M, 18F | 40.18 ± 10.10 | Ayahuasca | 3.6±0.2mg |
| Kiraga et al, 2021 | 19 | 14M, 5F | 39 ± 11.3 | Ayahuasca | 57.44 (±25.77) mg |

### Statistical Analyses

The meta-analyses were conducted using the ‘metaphor’ package [25] in RStudio version 2023.06.1 and Comprehensive Meta-Analysis (Version 3.3.070) [26]. Hedges’ g was used to measure effect size. Hedges’ g corrects for sample size bias and is interpreted according to Cohen’s D guidelines (0.2 for small effect, 0.5 for medium effect, and 0.8 for large effect). Q statistic was used to evaluate heterogeneity across studies based on the random effects model, as significant heterogeneity is expected in meta-analyses of observational studies [27]. Q assesses the amount of observed dispersion, and a statistically significant Q suggests that the studies do not share a common true effect size. The I2 index was used to assess the proportion of total variability in effect size estimates, with I2 values of approximately 25%, 50%, and 75% indicating low, moderate, and high heterogeneity, respectively. Publication bias due to small study bias was quantitatively tested using Egger’s regression test [28] for each outcome. An Egger’s test with a P value less than 0.05 indicated a small study bias. Outlier and influence diagnostics were conducted to determine the impact of any outliers and sensitivity re-analyses were performed without any identified outliers.

We gathered data for the MET task, which involved obtaining means and standard deviations for three variables: (1) cognitive empathy, which measured the accuracy (measured in percentage) in discerning emotions portrayed in the photos; (2) explicit emotional empathy, reflecting the participants’ ratings indicating their level of self-report explicit empathy towards the person in the photo; and (3) implicit emotional empathy, indicating the arousal levels experienced while viewing the photos.

## Results

### Classical Psychedelics effect on Cognitive empathy

Classical psychedelics did not affect cognitive empathy accuracy measures compared to placebo or baseline measures across the included studies (studies: n=5 combined overall sample size: n = 128, Hedges’ g=-0.1153 95% CI from -2.0316 to 1.8010, p=0.9061, Z-value=-0.1179). Between-studies heterogeneity was significant (Q = 36.7194, df = 5, I^2^ = 87.34%, p < 0.0001) (Figure 2).

**Figure 2.**
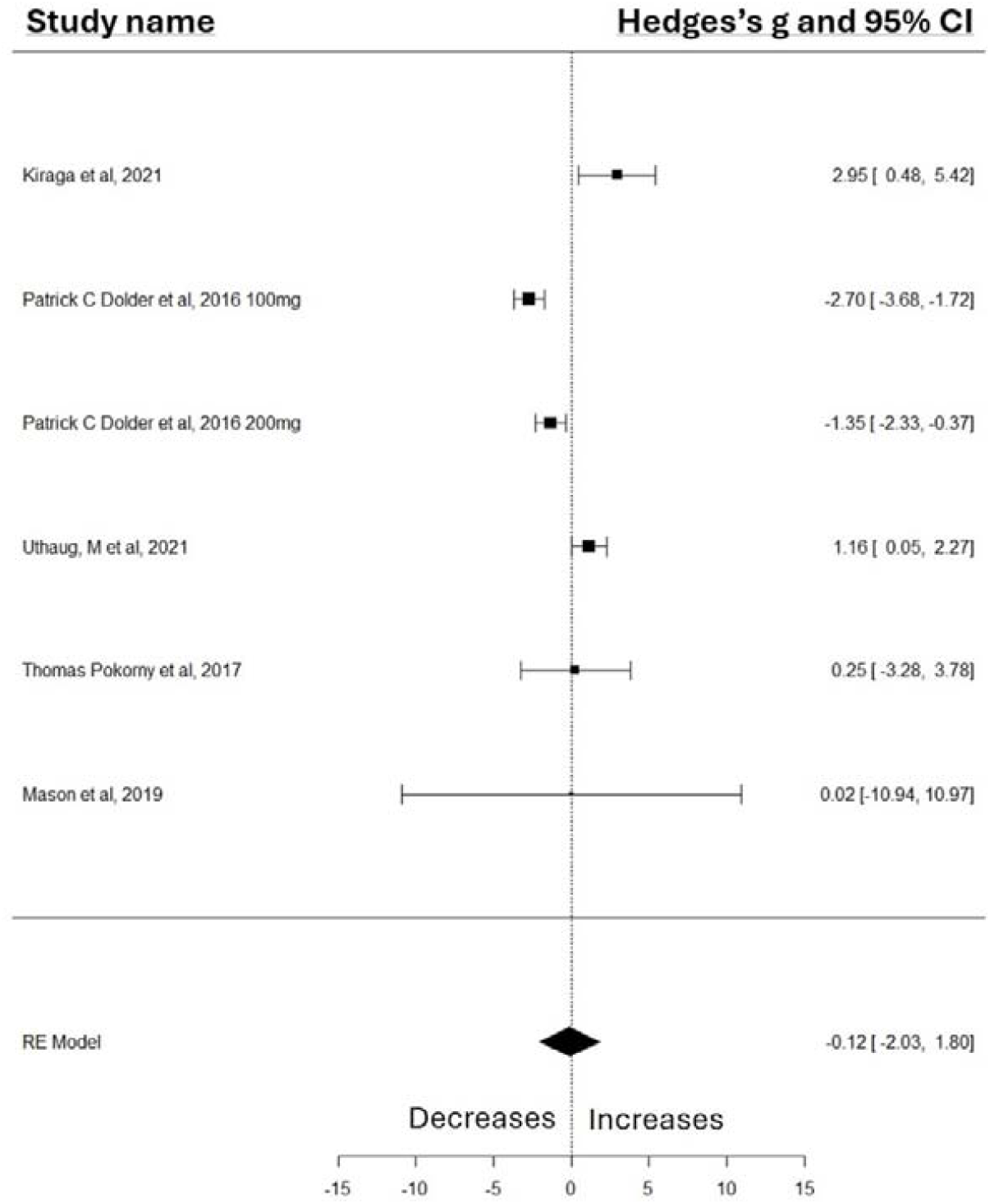
Forest Plot illustrates the influence of classical psychedelics intervention on cognitive empathy scores measured by the MET task.

### Publication Bias

No publication bias was found due to a small study effect, as indicated by Egger’s regression test (p = 0.5628). The funnel plot asymmetry tests, including regression test (z = 0.5786, p = 0.5628) and rank correlation test (Tau = 0.1380, p = 0.7021), did not indicate significant publication bias or small-study effects.

The trim-and-fill analysis, aiming to assess the potential impact of publication bias on the meta-analysis, revealed an estimated number of missing studies on the right side of 3, with a standard error (SE) of 1.5812. This suggests that the asymmetry observed in the funnel plot, indicative of potential publication bias, could have affected other studies. Influence diagnostics did not identify any outliers.

### Classical psychedelics’ Effect on Explicit Emotional empathy

Classical psychedelics were found to have a significant impact on explicit emotional empathy compared to placebo or baseline measures across the included studies (studies: n = 5; combined overall sample: n = 128, Hedges’ g = 0.7921, 95% CI from 0.1786 to 1.4057, p = 0.0114, Z-value = 2.5304). Between-studies heterogeneity was non-significant (Q = 1.1867, df = 5, I^2^ = 0.00%, p < 0.9461) (Figure 3).

**Figure 3.**
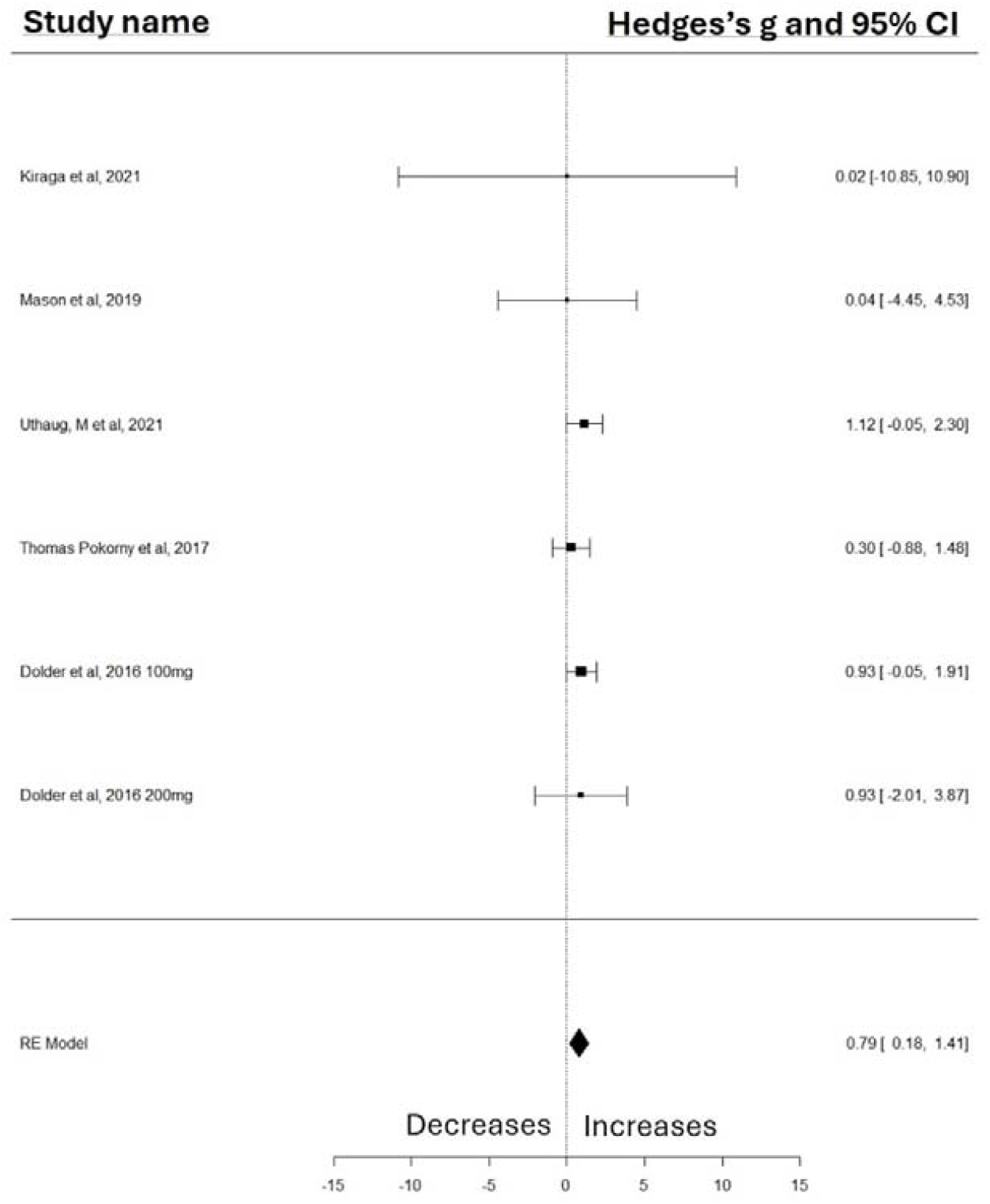
Forest Plot illustrates the influence of classical psychedelics intervention on explicit emotional empathy scores measured by the MET task.

### Publication Bias

There was no significant publication bias. This is evidenced by the absence of a small study effect, as indicated by Egger’s regression test (p = 0.7800). The results from the funnel plot asymmetry tests, including the regression test (z = -0.2793, p = 0.7800) and the rank correlation test (Kendall’s tau = -0.2760, p = 0.4442), consistently do not show significant publication bias or small-study effects. Influence diagnostics did not identify any outliers.

The trim-and-fill analysis, aiming to assess the potential impact of publication bias on the meta-analysis, revealed an estimated number of missing studies on the right side of 2, with a standard error (SE) of 1.7837. This suggests that the asymmetry observed in the funnel plot, indicative of potential publication bias, could have affected other studies. Influence diagnostics did not identify any outliers.

### Classical psychedelics effect on Implicit emotional empathy

Classical psychedelics were found to have a statistically significant impact on implicit emotional empathy ratings compared to placebo within the analyzed studies (studies: n = 5; combined overall sample: n = 128, Hedges’ g = 0.7635, 95% CI from -0.0191 to 1.5461, p = 0.05, Z-value = 1.9121).

The analysis revealed non-significant between-studies heterogeneity (Q = 0.4442, df = 5, I^2^ = 0.00%, p < 0.9940) (Figure 4).

**Figure 4.**
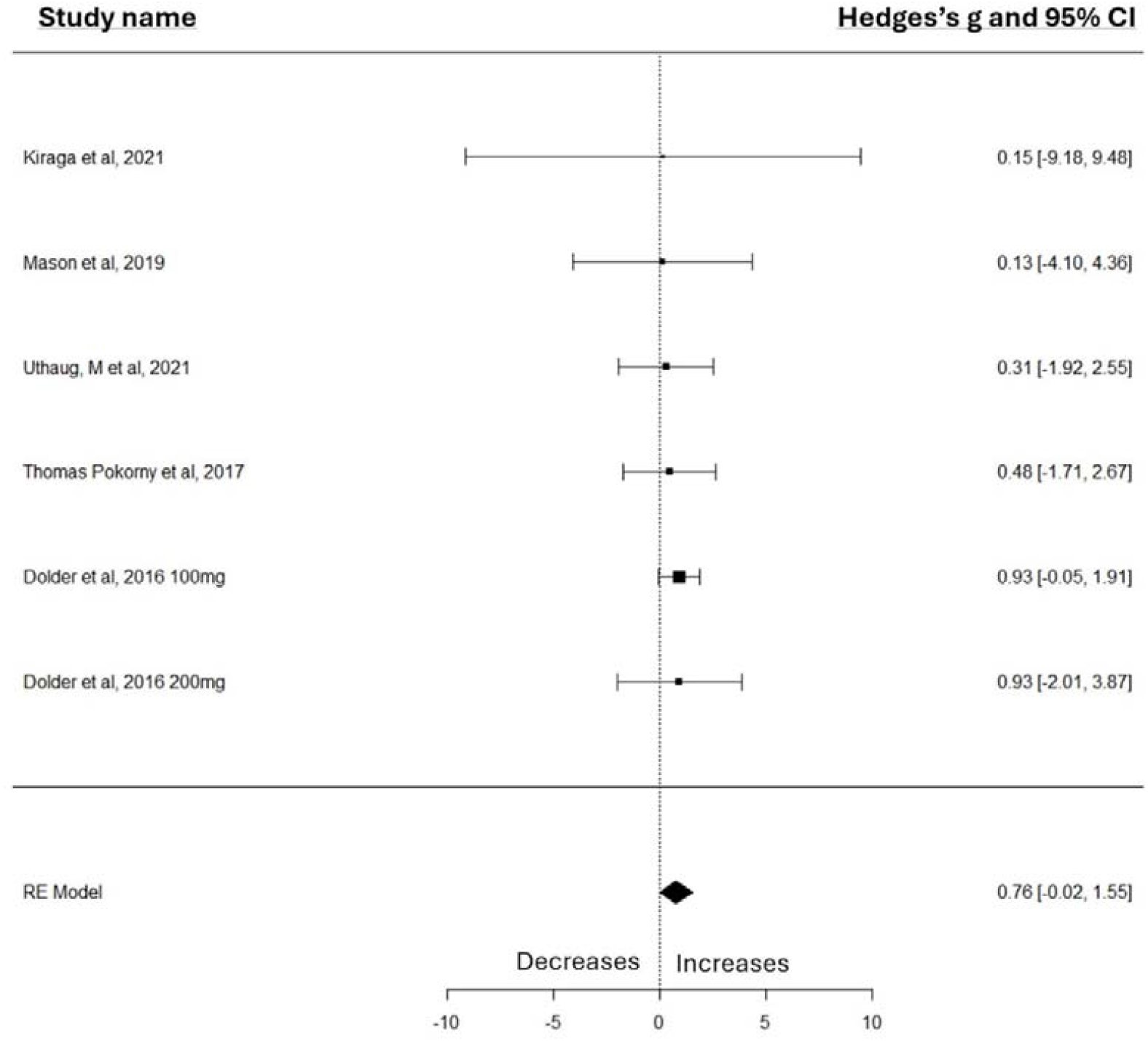
Forest Plot illustrates the influence of classical psychedelics intervention on implicit emotional empathy scores measured by the MET task.

### Publication Bias

Non significant publication bias was found, due to a small study effect as indicated by Egger’s regression test (p = 0.6427). The funnel plot asymmetry tests, including the regression test (z =-0.4639, p = 0.6427) and The Rank Correlation Test for Funnel Plot Asymmetry (Kendall’s tau of -0.2000 with a p-value of 0.7194), did not indicate significant publication bias or small-study effects.

The trim-and-fill analysis, aiming to assess the potential impact of publication bias on the meta-analysis, revealed an estimated number of missing studies on the right side of 3, with a standard error (SE) of 1.6385. This suggests that the asymmetry observed in the funnel plot, indicative of potential publication bias, could have affected other studies. Influence diagnostics did not identify any outliers.

## Discussion

The results of this meta-analysis indicate that classical psychedelics have a significant impact on the levels of explicit and implicit emotional empathy but no effect on cognitive empathy. The findings suggest that classic psychedelics may influence the emotional components of empathy, while further research is necessary to understand their impact on the cognitive components of empathy. There are several potential interpretations of these results. When using psychedelic substances, users have reported changes in perception [20], including hallucinations, subjective perception of time and space and perception of the body [29]. Specifically, distorted perception of the body might lead to blurring of the boundaries between self and other, which might lead to an increased ability to share the emotions of another, i.e., emotional empathy [4].

On a related note, psychedelic drugs can lead to a feeling of ego dissolution [17]. Ego dissolution entails the dismantling of the experience of the ego [17] and is accompanied by the lowering of the boundaries between the self and other people [10,17]. At the neural level, ego dissolution has been linked to reduced alpha power in the posterior cortex [30], which has been associated with increased emotional empathy levels [31,32]. It is possible that due to the effect of psychedelic drugs on the dissolution of the ego, there is an effect on emotional empathy, and this is due to an increase in the ability to merge with another person and, through that, to understand and share the feelings of others.

However, another mechanism through which classical psychedelics could affect emotional empathy is personality traits. Psychedelics were found to increase the personality trait of openness [33-36], and higher levels of openness have been associated with higher levels of general empathy [37]. Thus, it is possible that through an increase in openness to the experience of other individuals, the users can become more emotionally empathic to their surroundings. To get a deeper understanding of the relationship between psychedelics, empathy, and personality traits, further research is necessary.

A similar pattern of results was also observed with 3,4-methylenedioxymethamphetamine (MDMA), a compound with strong empathogenic properties [38-39]. Specifically, MDMA was also found to enhance emotional, but not cognitive, empathy [23,38]. Its action as a serotonin-enhancing agent shares common neurochemical pathways with classical psychedelics’ neural effects. This similarity suggests a potential overlap in how these substances modulate social and emotional processing, specifically through the enhancement of serotonin that affects emotional empathy but not cognitive empathy. Future research targeting serotonergic effects should determine the association between the enhancement of serotonin as a result of either MDMA or classical psychedelics and an increase in emotional empathy.

Our meta-analysis used a comprehensive and systematic approach to gather and analyze data. We examined the Multifaceted Empathy Test (MET), a standardized tool to measure empathy, across various studies. The use of the MET tool across all studies provided a consistent and specific measurement of empathy, enhancing the comparability and reliability of our results, which was the strength of this meta-analysis.

Moreover, our meta-analysis has a high ecological value since we took into consideration different settings and ways of administration of these substances. Specifically, the setting of a ceremony in cases of an ayahuasca ceremony and different physical, social and cultural environments in which the drug was taken may change the mental experience of their effect [40]. For example, one study reported that the ceremonies were held in a supportive group environment [10], which may affect the experience of the participants positively and may have contributed to improvements in the effect [41] compared to studies that examined the effects of the substance on individuals who were alone or with a therapist when they were administered the substance. Therefore, conducting a meta-analysis that analyzes results across these different settings contributes to understanding the isolated effect of these substances on empathy.

Moreover, the research includes different classical psychedelics, which enables us to generalize beyond one specific substance. The way psychedelics are reported in studies can be varied and complex due to differences in the types, doses, and timing of administration. In our research, the sample sizes ranged from 19 to 55) [11,19], ages from 25 to over 40 years [10,23], and dosage levels of psychedelics are different and depend on the type of drugs.

These results offer new possibilities for therapeutic applications. Empathy is crucial in psychotherapy because enhanced empathy can deepen the therapist-patient connection and improve therapy outcomes [9]. Specifically in disorders characterized by empathy deficits, such as certain personality disorders [20], psychedelics might aid in developing better social skills and emotional connections. Indeed, early studies suggested the use of psychedelic drugs as a treatment for criminals with antisocial personality disorders and psychopaths [42-44].

Importantly, we would like to acknowledge that all of the studies examined in this meta-analysis were done on healthy participants. Studying the effects of these substances on clinical populations could provide valuable insights into their therapeutic potential, such as depression and anxiety [14-16]. Moreover, to better understand the neurological effects of psychedelics, it is important to examine their acute and prolonged neural effect on empathy using imaging techniques such as electroencephalography (EEG) and functional magnetic resonance imaging (fMRI). These methods will provide insight into the neurological correlates of the empathy-enhancing effects of psychedelics.

Lastly, exploring individual differences in the subjective response to psychedelics and addressing ethical considerations surrounding their use are also crucial aspects of this field of research.

In conclusion, continued research on the effects of classical psychedelics on empathy is necessary for understanding their complex effects and potential therapeutic applications in fostering empathy and promoting prosocial behavior. Continued research in this domain is not just a scientific imperative but also a moral one, as it promises to look for new therapeutic modalities that could profoundly benefit individuals and society at large.

## Data Availability Statement

The datasets generated and/or analyzed during the current study are available from the corresponding author upon request. All data supporting this study’s findings are contained within the article and in supplementary information files.

